# Extracellular SPARC improves cardiomyocyte contraction during health and disease

**DOI:** 10.1101/492363

**Authors:** Sophie Deckx, Daniel M Johnson, Marieke Rienks, Paolo Carai, Elza Van Deel, Jolanda Van der Velden, Karin R Sipido, Stephane Heymans, Anna-Pia Papageorgiou

**Affiliations:** Department of Cardiology, Maastricht University, 6202 Maastricht, The Netherlands; Department of Cardiovascular Sciences, KU Leuven, 3000 Leuven, Belgium; Institute of Cardiovascular Sciences, University of Birmingham, Birmingham, UK; King’s British Heart Foundation Centre, King’s College London, London, UK; Department of Physiology, VUmc Amsterdam, 1081 Amsterdam, The Netherlands

**Keywords:** Cardiomyocyte, Contraction, Extracellular Matrix, SPARC, Viral Myocarditis

## Abstract

Secreted protein acidic and rich in cysteine (SPARC) is a non-structural extracellular matrix protein that regulates interactions between the matrix and neighboring cells. In the cardiovascular system, it is expressed by cardiac fibroblasts, endothelial cells, and in lower levels by ventricular cardiomyocytes. SPARC expression levels are increased upon myocardial injury and also during hypertrophy and fibrosis. We have previously shown that SPARC improves cardiac function after myocardial infarction by regulating post-synthetic procollagen processing, however whether SPARC directly affects cardiomyocyte contraction is still unknown. In this study we demonstrate a novel inotropic function for extracellular SPARC in the healthy heart as well as in the diseased state after myocarditis-induced cardiac dysfunction. We demonstrate SPARC presence on the cardiomyocyte membrane where it is co-localized with the integrin-beta1 and the integrin-linked kinase. Moreover, extracellular SPARC directly improves cardiomyocyte cell shortening *ex vivo* and cardiac function *in vivo, both* in healthy myocardium and during coxsackie virus-induced cardiac dysfunction. In conclusion, we demonstrate a novel inotropic function for SPARC in the heart, with a potential therapeutic application when myocyte contractile function is diminished such as that caused by a myocarditis-related cardiac injury.

## Introduction

In the heart, secreted protein acidic and rich in cysteine (SPARC) is expressed by endothelial cells, fibroblasts and in lower amounts by cardiomyocytes [1, 2]. SPARC is a collagen- and calcium-binding protein that belongs to the group of matricellular proteins. Matricellular proteins are matrix components characterized by (1) their counter-adhesive properties, (2) low expression levels during normal physiology but increased expression during stress and (3) the non-lethal phenotypes of knockout mice [2–4]. As typical of matricellular proteins, SPARC secretion occurs upon injury and at sites of remodeling in the heart. Previously, our group has shown that SPARC can improve clinical outcome after myocardial infarction by regulating the post-synthetic procollagen processing during fibrosis [5]. We showed how overexpression of SPARC could lead to an improved survival and increased cardiac contraction, as measured by echocardiography, after myocardial infarction in mice. Surprisingly, also in sham-operated mice, an increase in cardiac fractional shortening (FS) and ejection fraction was seen when SPARC was overexpressed. Yet whether SPARC directly affected cardiomyocyte contraction remained undetermined [5]. Therefore, the main aim of the present study was to investigate the potential inotropic function for SPARC in the healthy heart. Furthermore, as compromised cardiac contractile function is a hallmark of multiple cardiac diseases, we were also interested to investigate the therapeutic potential of SPARC in these conditions. For these reasons, a viral myocarditis (VM) model was utilized.

VM is an important inflammatory heart disease and an etiological precursor of dilated cardiomyopathy, (acute) heart failure and sudden cardiac death in young healthy individuals. Up to 60% of patients with dilated cardiomyopathy and myocarditis are virus-positive [6], yet diagnosis of VM is difficult due to its heterogeneous clinical presentation. Viral infection of the heart causes acute myocarditis, which can progress into chronic myocarditis causing cardiomyocyte damage and death, and initiation of remodeling processes such as fibrosis. All of these processes ultimately result in decreased contractile function of the heart as well as arrhythmia genesis and cardiac failure. Various viruses can cause viral myocarditis, including parvovirus B19, enteroviruses, hepatitis C virus and cytomegalovirus. The most studied are the coxsackie B viruses (CVB), which are members of the enteroviruses, and are often identified in biopsies from failing viral myocarditis hearts [7]. So far, research and development of novel therapeutic strategies for viral myocarditis has focused on processes targeting inflammation, cardiomyocyte degeneration, and fibrosis [7–9], whilst only a few studies have addressed the direct effect of viral infection on cardiomyocyte function. Importantly, viruses can also directly cause defective cardiomyocyte contraction, by time-dependently modulating numerous cardiac ion-channels, leading to alterations in action potential duration and resting membrane potential, as well as alterations in calcium loading which may contribute to viral-induced cardiac dysfunction [10–12]. Furthermore, non-structural matrix proteins in the heart can influence a myriad of processes during cardiac stress, such as inflammation, fibrosis and myocyte survival. Our group has previously demonstrated that the non-structural matrix proteins thrombospondin-2 and osteoglycin can affect inflammation, fibrosis and myocyte survival of the heart during cardiac aging, pressure overload, myocardial infarction, as well as after viral myocarditis [13–17] Recently we demonstrated that SPARC protects against adverse cardiac inflammation by preserving the endothelial glycocalyx during viral myocarditis [15]. Interestingly, we found clear differences in QTc times in SPARC KO mice as compared to WT mice during infection despite similar heart rates [15]. These data, in addition to our previous observation that SPARC overexpression leads to an increase in cardiac fractional shortening (FS) and ejection fraction, led us to investigate whether extracellular SPARC can act as an inotropic agent and influence cardiomyocyte contraction in health and during disease.

## Materials and Methods

### Mouse models

The Animal Care and Use Committee of the University of Leuven approved all described study protocols (ECD 243/2013). All animal studies conformed to the Guide for the Care and Use of Laboratory Animals. The Committee for Experiments on Animals of KU Leuven University, Belgium approved experiments. Animal handling was in accordance with the European Directive for the Protection of Vertebrate Animals used for Experimental and Other Scientific Purposes (2010/63/EU). For SPARC overexpressing experiments, an adenoviral vector designed by Barker *et al*. was used[18]. Adenovirus was produced by HEK293 cells that were collected and purified as previously described [19]. 1×10^10^ adenoviral PFU containing GFP or SPARC was injected into the tail of 12 week old C57Bl6 mice.

For viral myocarditis (VM) experiments, 3-5 week old male susceptible C3H mice (Harlan, Boxmeer, The Netherlands) were inoculated intraperitoneally with 10^3^ or 10^4^ PFU CVB3 (Nancy Strain) or PBS. Adenoviral overexpression experiments used the adenoviral vector designed by Barker *et al.* 1×10^10^ adenoviral PFU containing GFP or SPARC was injected into the tail vein of 3 weeks old mice 2 weeks prior to the CVB3 inoculation. For SPARC administration experiments, mice were subcutaneously infused for 72 hours with SPARC (40μg/kg/d) or vehicle (PBS) by Alzet osmotic minipump 1003D. Pump implantation surgery was performed as previously described [20] under ketamine and xylazine anesthesia at a dose of 100 mg/kg and 10 mg/kg respectively, and all efforts were made to minimize suffering. In all experiments, mice were sacrificed by a lethal injection of ketamine and xylazine (100mg/kg ketamine and 10mg/kg xylazine) intraperitoneally, plasma was collected, and hearts were removed and prepared for either myocyte isolation or histological and molecular analysis.

### Echocardiography analysis

Mice were anesthetized (2% isoflurane, ecuphar) and echocardiograpy was performed at indicated time points by transthoracic echocardiography with a 13-MHz transducer (i13L, GE ultrasound; Horton Norway) on a Vingmed Vivid 7 scanner (GE ultrasound, Horton, Norway). LV diameters at end-diastole (EDD), and end-systole (ESD), were measured, and fractional shortening (FS) was calculated.

### Adult mouse cardiac myocyte isolation and cell shortening experiments

Mice were injected with heparin (1000 U/kg intraperitoneally) and sacrificed by a lethal injection of ketamine and xylazine (100mg/kg ketamine and 10mg/kg xylazine) intraperitoneally. The heart was excised and cannulated via the aorta. Hearts were then mounted onto a Langendorff perfusion setup and initially briefly rinsed with normal Tyrode solution, containing (mM): 137 NaCl, 5.4 KCl, 0.5 MgCl2, 1 CaCl2, 11.8 Hepes, 10 2,3-Butanedione monoxime and 10 glucose, pH was adjusted to 7.4 with NaOH. Subsequently it was perfused with a Ca^2+^-free solution for 8 min. The Ca^2+^-free Tyrode solution contained (mM): 130 NaCl, 5.4 KCl, 1.2 KH2PO4, 1.2 MgSO4, 6 Hepes, 10 2,3-Butanedione monoxime, 20 glucose, and pH was adjusted to 7.2 with NaOH. Collagenase II (672Units/ml, Worthington 4176) added to the Ca^2+^-free solution was subsequently perfused for 8 min. The enzyme was then washed out for a further 3 mins with 0.09mM CaCl2 and 50mg/ml BSA 0.18 mM CaCl2. The heart was then removed from the Langendorff perfusion setup, and the myocytes were further dissociated mechanically by gentle shaking. Ca^2+^ was reintroduced stepwise.

Cell shortening was measured using video-edge detection (Ionoptix) during electrical field stimulation at 1 and 2 Hz. Field stimulation was achieved with 5 ms square pulses of constant voltage, at 20 *%* above threshold. The cell shortening is expressed as the fractional shortening, i.e. normalized to resting cell length, AL/L0*100%. During field stimulation cells were superfused with normal Tyrode solution at 37°C. [Ca2+]i was measured with fluo-3, and is reported as the fluorescence normalized to baseline values, after background subtraction, F/F0. To measure the effect of SPARC *ex vivo* on cells, recombinant SPARC (1μg/ml) was added to half of the freshly isolated cell suspension, while the other half was left in normal Tyrode solution. Additionally, to assess the effects of integrin-linked kinase (ILK) inhibition, CPD-22 (Calbiochem; 1μM) was utilized, whilst to investigate the effect of myosin light chain kinase (MLCK) inhibition, ML-7 was used (Sigma Aldrich; 3 μM). For all of the above interventions myocytes were incubated with the specific agent(s) for 1 hour and then cell shortening was measured as described above.

### In vitro experiments with adult rat cardiac myocytes

Cardiac myocytes were isolated by enzymatic dissociation from adult Wistar rat hearts as previously described [21]. Experiments were performed in accordance with the Guide for the Animal Care and Use Committee of the VU University Medical Center (VUmc) and with approval of the Animal Care Committee of the VUmc. For experiments, freshly isolated cardiomyocytes were cultured at 37°C overnight on polyarcrylamide gels (25% Acrylamide40%, 13% Bis2%) with a stiffness of approximately 15kPa. Prior to culture, these gels were coated with laminin (10μg/ml), collagen (50μg/ml) with and without recombinant SPARC (1μg/ml, PeproTech 120-36) in 0.1M HEPES, overnight at 4°C. The cardiac myocytes were plated in plating medium (M199 medium, Gibco, 31150-022, with 1% Peniciline-Streptavidine and 5% fetal bovine serum) onto the coated gels, and after 1 hour incubation at 37°C, medium was replaced to culture medium (M199 medium with 1% Peniciline-Streptavidine, 0.2% Insulin Transferrin Sodium selenite and 0.1% Cytochalasin D). After overnight incubation at 37°C, unloaded cell shortenings of the adherent cardiac myocytes were measured in the culture medium, using different frequencies of electrical field stimulation and analyzed using IonOptix software (IonOptix LLC, Milton, MA). Data are presented as fractional shortening (% diastolic length), time to peak of contraction (TTP) and 50% relaxation time (RT50).

### Determining gel stiffness

Stiffness of fully hydrated gels was determined using a Piuma Nano-indentor (Optics 11, Amsterdam, the Netherlands) in combination with an indentation probe with a stiffness of 1 N/m and a tip radius of 44 um (Optics 11, Amsterdam, the Netherlands). The gel’s Young’s modulus was determined by averaging 9 individual measurements.

### Histology and microscopy

Cardiac tissue was processed and histochemical and immunohistochemical analyses were performed as previously described [22–24], and all morphometric analyses were done on sections with myocyte in cross-sectional images. Hematoxylin and eosin – stained sections (4 μm) were used to assess overall morphology. The number of CD-45 – staining cells (monoclonal rat antibody, BD, 553076, clone 30-F11, 5μg/ml) was measured per mm^2^. Myocyte cross-sectional areas were calculated by measuring the inner circumference of 150 myocytes per sample on laminin–stained sections (rabbit antibody, Sigma, L9393, 125μg/ml). To assess the amount and cross-linking of fibrosis, Picro Sirius Red staining was performed as previously described [24, 25]. Microscopic analyses were performed using a microscope (Leitz DMRXE; Leica), and QWin morphometry software (Leica). All analyses were performed according to standard operating procedures.

### Immunostaining of isolated cardiac myocytes

Adult cardiac myocytes were isolated from healthy mice as described, fixed in 2% PFA in PBS for 10 min, incubated in 50mM glycine for 30 min to remove auto-fluorescence caused by PFA at 488nm, and subsequently stained for SPARC (polyclonal goat antibody, R&D systems, AF492, 5μg/ml) overnight at 4°C. The next day cells were initially incubated with a secondary donkey-anti goat-alexa 488 labeled antibody for 90min. at room temperature and some cells were subsequently stained for integrin beta1 (monoclonal rat antibody, BD, 553715, 0.5μg/ml) for 4 hours at room temperature and afterwards incubated with a secondary goat-anti rat-alexa 568 labeled antibody for 90min. at room temperature. Cells were visualized with confocal microscopy on a Zeiss LSM700 microscope (Leica) using the Zen software (Leica), or analyzed using a BD FACSAria III flow cytometer (Becton Dickinson (BD), San Jose, CA) and FlowJo software (Ashland, Oregon).

### Immunostaining of the coated matrices

Matrices were produced and coated as described previously and stored at 4°C prior to staining. Matrices were washed with PBS and subsequently stained for SPARC (polyclonal goat antibody, R&D systems, AF491, 5μg/ml), laminin (polyclonal rabbit antibody, Sigma, L9393, 5μg/ml) and collagen (monoclonal rat antibody, Merck Millipore, MAB 1912, 1/100) overnight at 4°C. The next day matrices were incubated with a secondary donkey-anti goat-alexa 660, goat-anti rabbit-alexa 568 and, goat-anti rat-alexa 488 labeled antibodies for 90min. at room temperature and matrices were visualized with confocal microscopy on a Zeiss LSM700 microscope (Leica) using the Zen software (Leica).

### Myocyte fractionation

Adult cardiac myocytes were isolated from healthy mice as described, incubated in lysis buffer, containing (mM): 5 TrisHCl, 5 NaCl, 2 EDTA, 1 CaCl2, 1MgCl2, 2 DTT and pH was adjusted to 7.4. Phosphatase inhibitors (2%, Sigma, P044 and P5726) and protease inhibitors (4%, Roche, 11697498001) were added to the buffer, and cells were incubated overnight at 4°C. The next day, the cell suspension was centrifuged for 1 hour at 4°C and supernatant was collected as cytoplasmic fraction, the pellet was dissolved in lysis buffer and collected as the membrane fraction.

### Immunoprecipitation

For immunoprecipitation, left ventricular tissue or isolated cardiomyocytes were lysed in immunoprecipitating buffer containing (mM): 150 NaCl, 20 Tris, 5 EDTA, 1% Triton X-100 and pH was adjusted to pH 7.5 using NaOH. Phosphatase inhibitors (2%, Sigma, P044 and P5726) and protease inhibitors (4%, Roche, 11697498001) were added to the buffer. Dynabeads M-280 (Sheep anti rabbit a-Ig, Life Technologies, 2018-06) were washed with lysis buffer and incubated with SPARC antibody (monoclonal rabbit antibody, Sino Biological Inc, 50494-R001, 3ug in 200uL buffer) or rabbit serum as negative control, for 2 hours at 4°C. Next, beads were washed and incubated with lysates overnight at 4°C. The next day, the non-bound lysates were collected and resolved for SDS-PAGE, beads were washed and beads-bound immune complexes were resolved for SDS-PAGE. Samples were subsequently immunoblotted for the detection of ILK (polyclonal rabbit antibody, CST, 3862, 1/1000).

### Western Blotting

Proteins were isolated from left ventricular tissue, or from isolated cardiomyocytes, separated by SDS-PAGE and subsequently immunoblotted for the detection of pAkt (monoclonal rabbit antibody, Cell signaling, 4060, 1/1000), and total Akt (polyclonal rabbit antibody, Cell signaling, 9272, 1/1000), SPARC (polyclonal goat antibody, R&D systems, AF492, 5μg/ml) and GAPDH (monoclonal mouse antibody, Fitzgerald, 10R-G109a, clone 6C5, 0.1μg/ml) overnight at 4°C. Signals were visualized using Hyperfilm ECL (Amersham Biosciences) and quantified using Image J software.

### Statistics

Data were expressed as the mean ± SEM. Histological and molecular analyses in sham-operated and VM groups were performed in independent groups. For echocardiographic measurements, analyses were performed in independent groups, except for the experiment where SPARC or vehicle was infused with an osmotic minipump for 72h, where repeated measures were performed. Normal distribution of all continuous variables was tested according to the method of Kolmogorov and Smirnov. An unpaired Student t-test for 2 groups or ANOVA, followed by a Bonferroni post hoc test for more groups was used in most of the comparisons when groups passed the normality test.

When the standard deviation of two groups significantly differed, a Mann-Whitney test for 2 groups or a Kruskal-Wallis test, followed by a Dunn’s post hoc test for more groups, was used. A paired Student’s t test was used to analyze baseline and follow-up echocardiographic measurements, a Wilcoxon test was used when data did not pass normality test. A two-sided p-value of ≤ 0.05 was considered statistically significant.

## Results

### Extracellular SPARC increases cardiomyocyte contraction

To study how extracellular SPARC can improve cardiac function, we investigated whether extracellular SPARC directly interacts with the cardiomyocytes. SPARC presence on the cardiomyocyte increased when cells were isolated and incubated with 1mg/ml SPARC ex vivo for 1h, as compared to cells incubated in normal buffer without SPARC (Figure 1A). These SPARC-incubated cells demonstrated a higher cardiomyocyte cell shortening (Figure 1B and C), with no significant changes in contraction times (TTP) or relaxation times (RT50) (Supplementary Figure 1A and B). To mimic more *in vivo* conditions, we coated matrices with physiological stiffness with laminin and collagen (Lam+Col) or laminin, collagen and SPARC (Lam+Col+SPARC) (Figure 1G). SPARC, a known collagen-binding protein, co-localized with collagen on the matrices (Figure 1D) and importantly, the presence of SPARC did not alter matrix stiffness (Figure 1E). Next, we isolated cardiomyocytes from adult rats and cultured them overnight on these matrices. When stimulated at 0.5, 1 and 2 Hz, rat cardiomyocytes cultured on SPARC-containing matrices demonstrated an increased cardiomyocyte shortening at all frequencies when compared to cells cultured on matrices without SPARC (Figure 1F), while, once again, both TTP and RT50 were unchanged (Supplementary Figure 1C and D).

**Figure 1.**
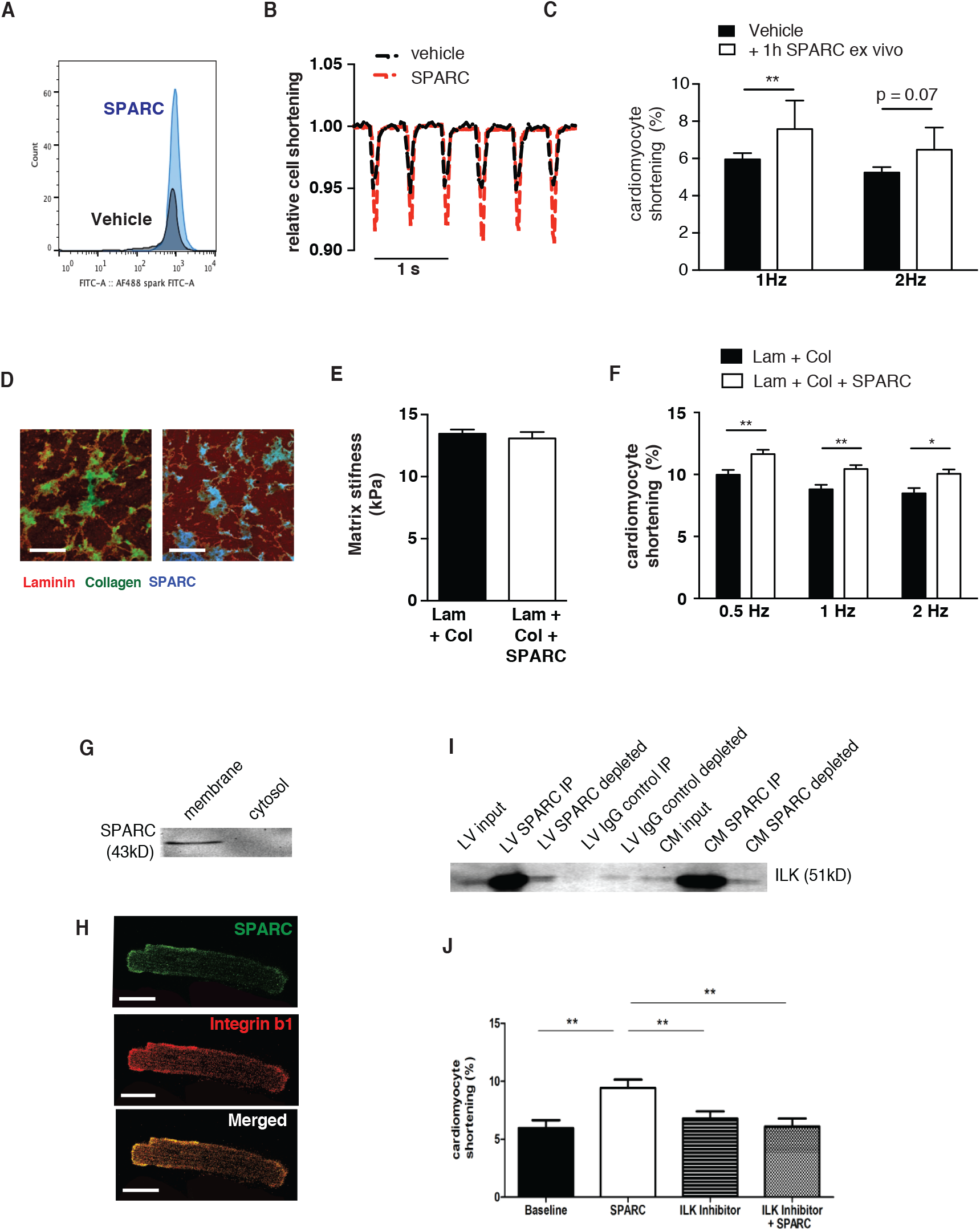
SPARC improves cardiomyocyte contraction through its interaction with integrin-beta1. A FACS analysis demonstrates increased SPARC staining when cardiomyocytes were isolated and incubated with SPARC *ex vivo* for 1h, as compared to cells incubated in normal buffer without SPARC. B, C Isolated adult mouse cardiomyocytes displayed higher FS after 1h incubation with SPARC, compared to cells incubated in normal tyrode buffer. D, E Matrices with physiological stiffness were coated with laminin and collagen (Lam + Col) or laminin, collagen and SPARC (Lam + Col + SPARC). SPARC co-localized with collagen on these matrices, but did not affect matrix stiffness. F Adult rat cardiomyocytes were isolated and cultured on these matrices. Cells cultured on SPARC containing matrices displayed higher fractional shortening (FS), compared to cells cultured on matrices coated with L+C alone. G SPARC is present in the membrane fraction, and absent in the cytosolic fraction of isolated cardiomyocytes, as demonstrated by Western Blotting. H Using immunostaining and confocal microscopy we confirmed SPARC presence on the cardiomyocyte membrane, where it colocalizes with integrin-beta1. I SPARC immunoprecipitation (I.P.) demonstrates interaction with integrin-linked kinase (ILK) in LV samples and in isolated cardiomyoctes. J Isolated adult mouse cardiomyocytes were incubated in the presence of SPARC and/or the ILK-inhibitor CPD-22. The increased FS observed in cells incubated with SPARC was abolished in the presence of CPD-22. A-C N=4 mice and n >4 cells per mouse, D-F N= 3 rats and >20 cells per rat, J N = 3 rats and >4cells per rat, bars panel D 100um, bars panel H 10um, *p<0.05, **p<0.01

Using Western Blot, we demonstrate SPARC presence in the membrane fraction, yet absence in the cytosolic fraction of isolated cardiomyocytes (Figure 1G). Using immunostaining and confocal microscopy we confirmed SPARC’s presence on the membrane of the cardiac myocyte, where it co-localizes with integrin-beta1 (Figure 1H). Furthermore, immuno-precipitation demonstrated an interaction of SPARC with integrin-linked kinase (ILK) in both whole LV samples and in isolated cardiomyocytes (Figure 1I).

To investigate whether this SPARC-induced increased cardiomyocyte cell shortening is through ILK-signaling, we incubated cells in the presence of SPARC and/or the ILK-inhibitor CPD-22. Importantly, the SPARC-induced increased cardiomyocyte cell shortening is blunted in the presence of the ILK-inhibitor (Figure 1J). These results indicate that SPARC increases cardiomyocyte cell shortening, at least in part, through ILK signaling. Notably, CPD-22 alone did not affect cardiomyocyte shortening (Figure 1J).

In conclusion, these results demonstrate a direct binding of SPARC with the cardiomyocyte membrane, where it appears to interact with ILK and is found in close proximity to integrin-beta1. Moreover, SPARC presence on the membrane increases when cells are incubated in the presence of recombinant SPARC, resulting in increased cardiomyocyte contraction, mediated -at least in part-through increased ILK signaling.

### SPARC improves cardiomyocyte function in virus-induced heart failure

We subsequently investigated whether SPARC was able to improve cardiomyocyte fractional shortening in conditions where function of this cell type is impaired. Considering the influence of SPARC on enhancing collagen cross-linking, we aimed to study the influence of SPARC on cardiomyocyte function *in vivo* in a disease model with limited fibrosis. As we recently observed changes in cardiac function and ventricular conductivity during VM when SPARC was absent, we decided to investigate in further details the effect of SPARC on cardiomyocyte function in this disease setting. Therefore, a low-dose VM model where mice developed minimal fibrosis, allowed us to study the effect of adenoviral mediated SPARC overexpression on cardiac function and cardiomyocyte contraction. In this model 103 PFU CVB3 was injected intraperitoneally, resulting in mild inflammation and fibrosis, and no cardiomyocyte hypertrophy (Figure 2A – D). Yet, cardiac contraction as measured by FS was decreased (Figure 2E), and there was an apparent onset of cardiac dilation suggested by an increased end systolic diameter (ESD) (Figure 2F). Importantly, fractional shortening of isolated cardiomyocytes was also compromised in this model (Figure 2G). Under field stimulation conditions, these cardiomyocytes did not display a prolonged TTP, but RT50 values were significantly increased (Supplementary Figure 2A and B).

**Figure 2.**
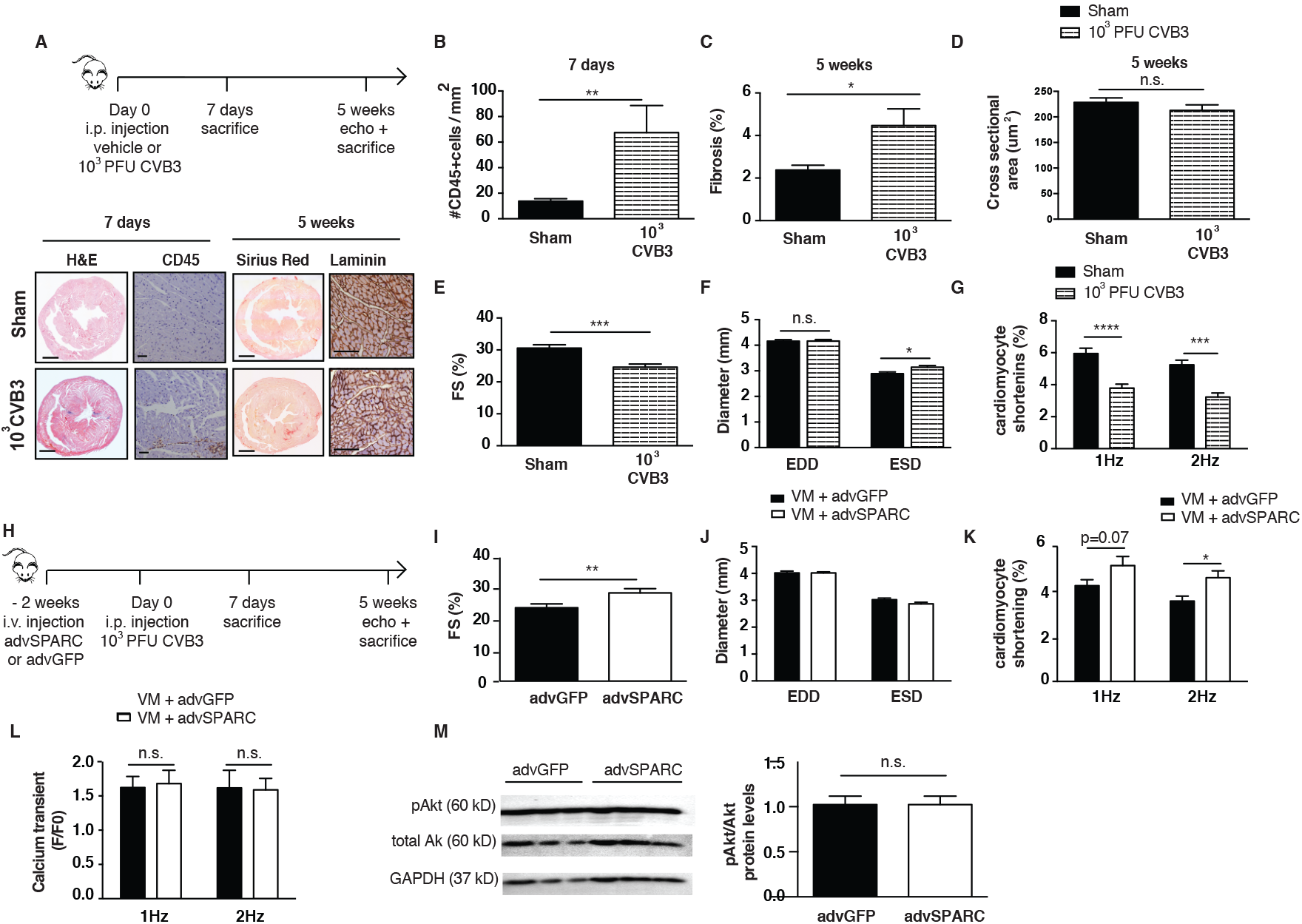
SPARC improves cardiomyocyte function in a mild model of virus-induced heart failure. A – D A mild VM mouse models is used, where mice are injected with10^3^ PFU CVB3 intraperitoneally. This results in moderate cardiac inflammation after 1 week), little fibrosis and no cardiomyocyte hypertrophy after 5 weeks. E,F Viral infection caused decreased FS and increased ESD. G Contraction of isolated cardiomyocytes is also compromised after virus-infection. H SPARC is overexpressed with the use of an adenovirus, 2 weeks before mild CVB3 inoculation. I,J 5 weeks after CVB3 injection, higher FS were measured in the SPARC overexpressing group, with no differences in EDD, and slightly smaller ESD. K Isolated myocytes from the SPARC-overexpressing hearts remained their increased shortening capacities as compared to isolated myocytes from control GFP-hearts. L,M There were no differences in the Ca^2+^ transient peak heights, or levels of Akt phosphorylation. A-F n= 11 for sham and n=13 for VM, G n= 11 for sham and n=13 for VM and >3 cells per mouse, H-J n=12 for advGFP group and n= 11 for advSPARC group, K,L n=12 for advGFP group and n= 11 for advSPARC group and >3cells per mouse, M n=3 for both groups, bar 1000um for H&E and Sirius Red stainings, 100um for CD45 and laminin stainings, *p<0.05, **p<0.01, ***p<0.001, ****p<0.0001

Next, using this low-dose VM model, we systemically injected SPARC with the adenoviral vector with the intention of increasing cardiac SPARC expression (Figure 2H). One week after CVB3 injection, Western Blotting revealed that cardiac SPARC levels were increased in 3 out of the 4 mice from the adenoviral-SPARC injected group when compared to the control adenoviral-GFP injected mice (Supplementary Figure 2C). Five weeks after CVB3 injection, higher FS, and preserved ESD was measured in the SPARC overexpressing animals as compared to GFP overexpression (Figure 2I,J), demonstrating that SPARC overexpression prevents the development of cardiac dysfunction in this mild VM model. Importantly, myocyte cross-sectional area, the amount of fibrosis, collagen cross-linking, and the number of CD45 positive cells in the heart did not differ between the 2 groups, 5 weeks after CVB3 injection (Table 1). Still, increased fractional shortening was demonstrated by isolated cardiomyocytes from SPARC-overexpressing animals as compared to isolated cardiomyocytes from control GFP overexpressing animals (Figure 2K), indicating a protective, or positive inotropic effect of SPARC at the level of the cardiomyocyte. Notably, no effect on contraction or relaxation times was observed (Supplementary Figure 2E and F). Furthermore, despite SPARC being a Ca^2+^-binding protein, we could not find indications that SPARC influenced Ca^2+^-handling, as there were no differences in the Ca^2+^ transient peak heights (Figure 2L), TTP or RT50 (Supplementary Figure 3G and H) of these isolated myocytes. Moreover, we did not find any differences in Akt phosphorylation between LV samples from both groups, which is known to increase intracellular Ca^2+^-availability and enhance contraction [26], in LV samples from both groups, as shown by Western Blotting (Figure 2M), further supporting no immediate role for SPARC in Ca^2+^-handling. Taken together, these data demonstrate a protective effect of SPARC on cardiomyocyte function prior to the establishment of virus-induced heart failure. Furthermore, these data indicate that SPARC affects filament sensitivity to Ca^2+^ rather than altering Ca^2+^ handling within the cell.

**Table 1.** Histological analysis of VM mice with GFP or SPARC adenoviral overexpression

|  | 5 weeks VM |  |
| --- | --- | --- |
|  | AdvGFP<br>(n=10) | AdvSPARC<br>(n=8) |
| **Myocyte cross-sectional area ($\mu\text{m}^2$)** | 217± 6 | 224 ± 7 |
| **Fibrosis (%)** | 2.4 ± 0.2 | 2.6 ± 0.4 |
| **Orange-red/ yellow-green fibers** | 1.15 ± 0.17 | 1.61 ± 0.14 |
| **CD45+ cells / mm<sup>2</sup>** | 6.67 ± 2.73 | 6.57 ± 3.0 |

**Table 2.**
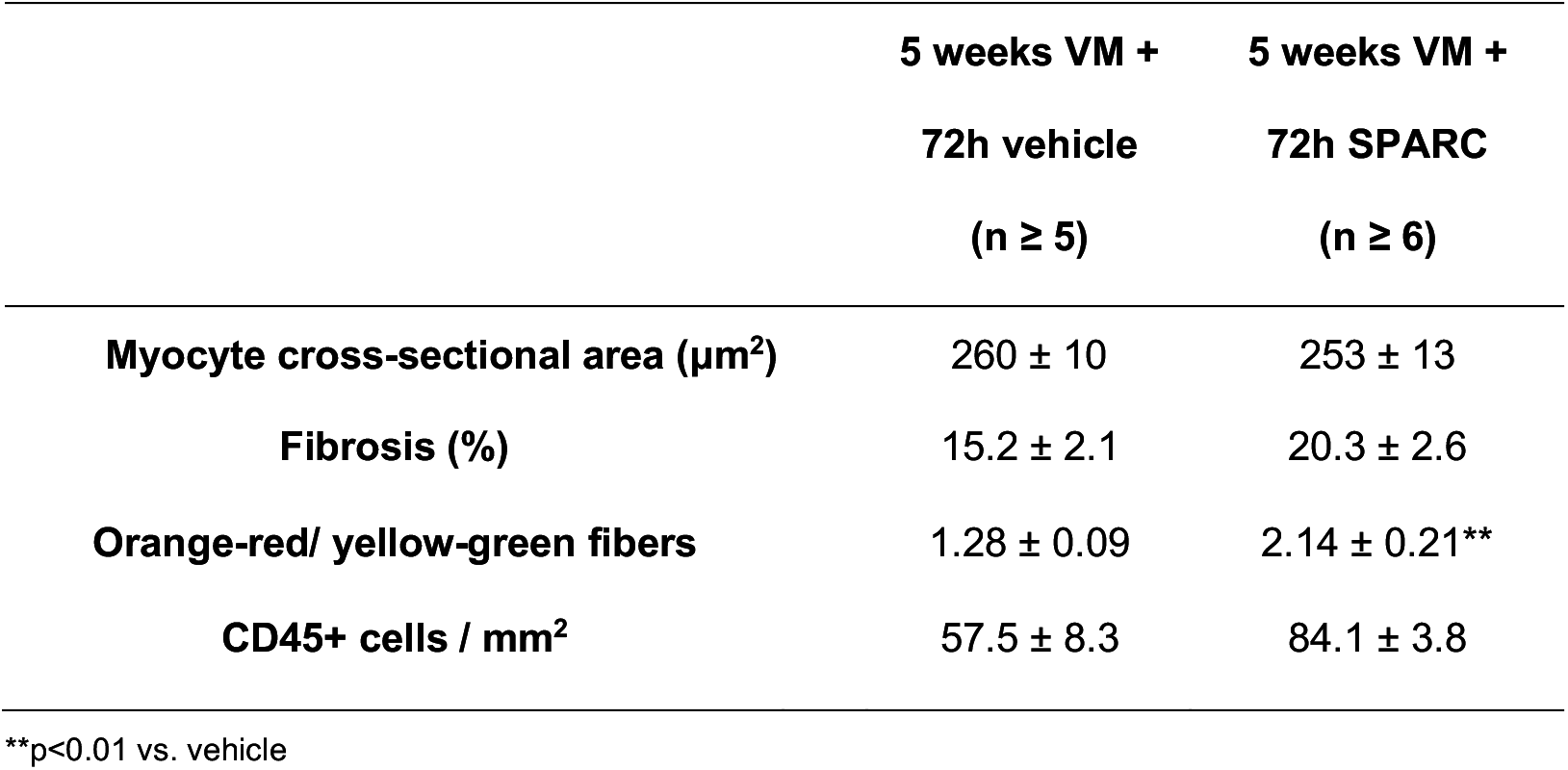
Histological analysis of VM mice with 72h vehicle or SPARC infusion

We next wanted to assess the therapeutic potential for SPARC, using a high-dose CVB3 model with pronounced cardiac inflammation and fibrosis and severely compromised cardiac function (Figure 3A-F). In this model, a higher dose of CVB3 (10^4^ PFU CVB3) was injected intraperitoneally in mice, resulting in severe cardiac inflammation after 1 week, and prominent fibrosis and cardiomyocyte hypertrophy after 5 weeks (Figure 3A – D). Here, cardiac function was even more compromised, as shown by severely decreased FS (Figure 3E), and significantly increased ESD, indicating cardiac dilation in this model (Figure 3F). To investigate the therapeutic potential of SPARC, we infused mice with SPARC or vehicle for 72h implanting an osmotic minipump 5 weeks after initial viral exposure, when dilated cardiomyopathy with severe inflammation and fibrosis had been established and measured cardiac function prior and after 72h of SPARC or vehicle infusion (Figure 3G). We found an increased FS in the SPARC treated group, while FS in the vehicle group continued to decline (Figure 3H). End diastolic diameter (EDD) was slightly smaller in the SPARC group prior to treatment, compared to the vehicle group.

**Figure 3.**
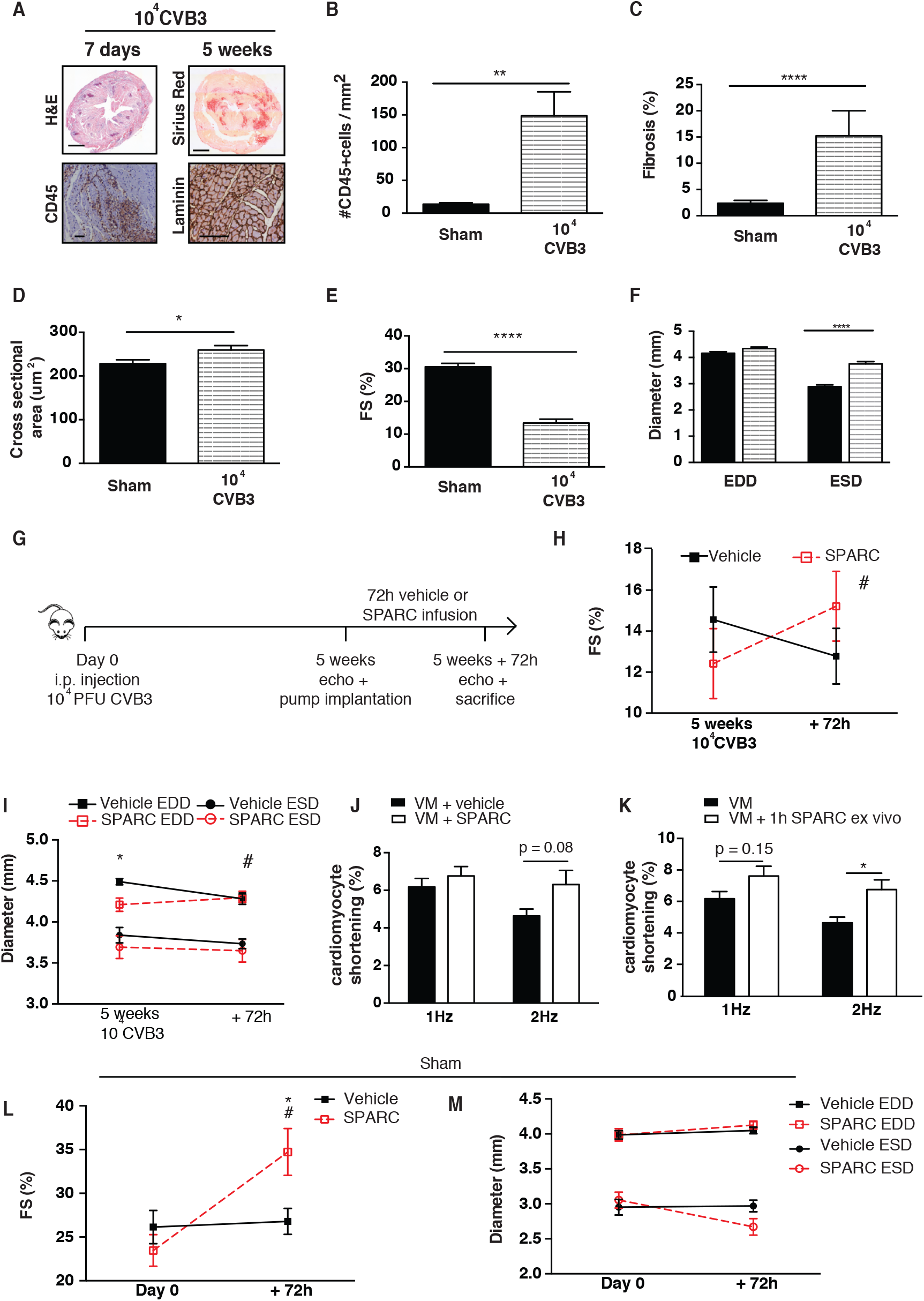
SPARC has therapeutic potential in severely virus-induced heart failure. A – D In the more severe VM model mice are injected with 10^4^ PFU CVB3, which results in severe cardiac inflammation after 1 week, and prominent fibrosis, but no cardiomyocyte hypertrophy after 5 weeks. E, F Viral infection caused severely decreased FS and dilation of the heart. G Mice were infused with SPARC or vehicle for 72h, 5 weeks after high-dose CVB3 inoculation, when dilated cardiomyopathy with severe inflammation and fibrosis had been established. H FS was increased in the SPARC treated group, while FS in the vehicle group continued to decline. I EDD were slightly smaller in the SPARC group prior to treatment, compared to the vehicle group, but did not change due to the SPARC treatment, while in the vehicle group EDD were slightly decreased after 72h. ESD were not different between groups or between time-points. J FS was increased in isolated cardiomyocytes from SPARC-treated mice when compared to cells isolated from vehicle-treated mice. K When cardiomyocytes were isolated from the severely sick, untreated mice, incubation of the cells with SPARC for 1h *ex vivo* also resulted in increased FS, compared to control cells. L,M Also healthy mice demonstrated higher FS when SPARC was administered for 72 hours, compared to vehicle-administered mice. This resulted in decreased ESD but not decreased end-diastolic diameters EDD in SPARC-administered mice, while diameters did not change in vehicle-administered mice. A-F n=11 for sham and n=13 for VM, G-I n=6 for VM+vehicle and n=7 for VM+SPARC, J n=6 for VM+vehicle and n=7 for VM+SPARC and >3cells per mouse, K n=13 for both groups and >3cells per mouse, L,M n=11 for sham+vehicle and n=8 for sham+SPARC, bar 1000um for H&E and Sirius Red stainings, 100um for CD45 and laminin stainings, *p<0.05, **p<0.01, ***p<0.001, ****p<0.0001

However, EDD did not change due to the SPARC treatment, while in the vehicle group EDD were slightly decreased after 72h. ESD, on the other hand, was not different between groups or time-points (Figure 3I). Moreover, myocyte cross-sectional area and the amount of CD45 positive cells in hearts did not differ between the 2 groups (Table 2). Yet, while the amount of fibrosis did not differ, collagen cross-linking was increased in the SPARC-treated group as compared to the vehicle group (Table 2), confirming the previously demonstrated effect of SPARC on collagen-crosslinking. Nevertheless, despite this higher collagen cross-linking, we found next to increased cardiac contraction, a trend to increasing shortening in isolated cardiomyocytes from these SPARC-treated mice at 2Hz pacing cycle length when compared to cells isolated from vehicle-treated mice (Figure 3J), with no differences in TTP or RT50 (Supplementary Figure 2I and J). In addition, when cardiomyocytes were isolated from these severely sick, untreated mice, incubation of the cells with SPARC for 1h *ex vivo* resulted in a significant increase in cardiomyocyte shortening, compared to control cells (Figure 3K), again without influencing TTP or RT50 (Supplementary Figure 2K and L).

Finally, when we infused healthy adult mice with SPARC or vehicle for 72h, we also found increased FS compared to baseline measurements and compared to vehicle-mice (Figure 3L). SPARC administration caused decreased end-systolic diameters (ESD), but not end-diastolic diameters (EDD), while diameters did not change in hearts of vehicle-mice (Figure 3M) Again, SPARC administration did not affect cardiomyocyte hypertrophy, the amount of fibrosis, collagen cross-linking, or the amount of CD45 cells (Table 3)

**Table 3.**
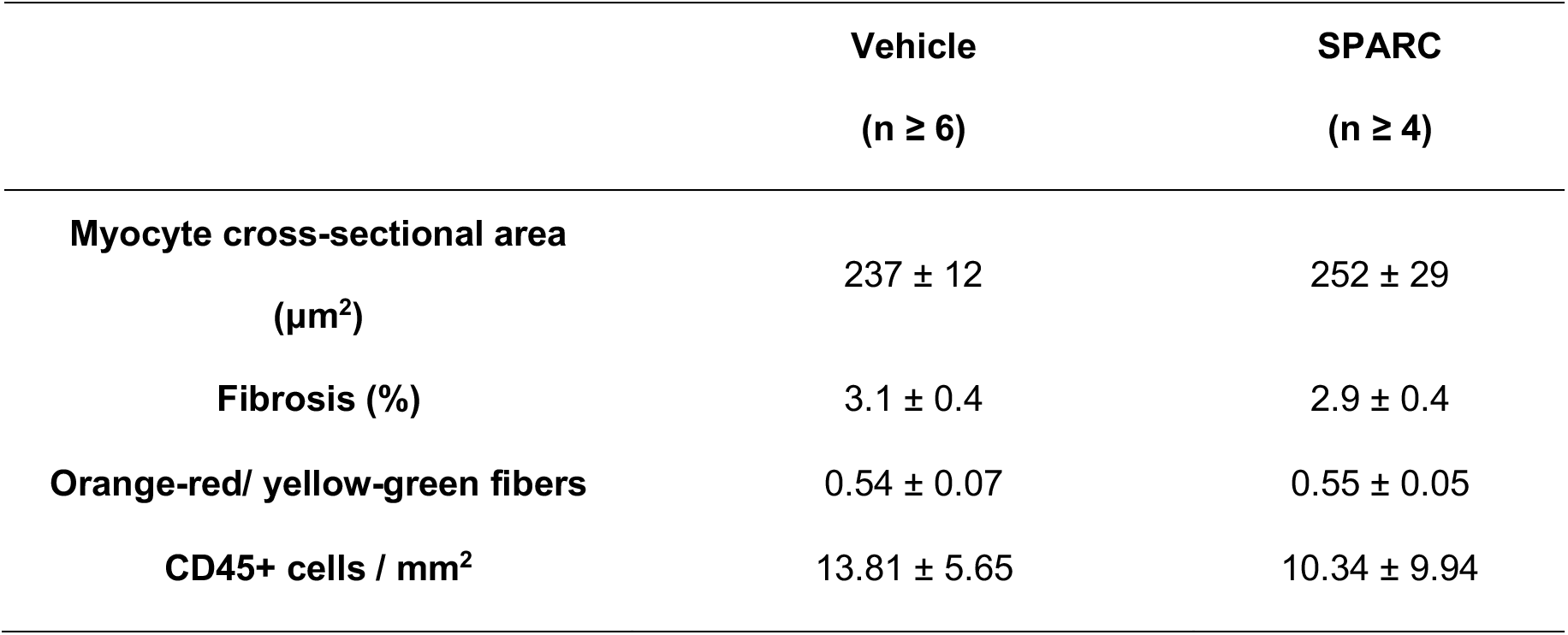
Histological analysis of hearts after 72h vehicle or SPARC administration

## Discussion

To our knowledge this study is the first to demonstrate a direct role for a non-structural matrix protein on cardiomyocyte contraction, via interactions of this protein with intracellular effectors. Our previous study on SPARC in myocardial infarction suggested a previously unexplored potential inotropic function for SPARC in the heart. Here, we demonstrated increased cardiac contraction when SPARC was overexpressed, not only in infarcted mice, but also in sham-operated mice [5] yet how SPARC might directly affected cardiomyocyte contraction remained undetermined. Therefore, in this study we aimed to explore the role of SPARC on cardiomyocyte contractility using various *ex vivo* and *in vivo* models. We have shown that extracellular SPARC increases cardiomyocyte contraction, during both health and disease, possibly by interacting with the integrin-beta1-ILK complex on the cardiomyocyte membrane. Not only is SPARC able to prevent a decrease in cardiac function, but it is also able to rescue myocytes that are already compromised through viral infection. These data highlight the potential of SPARC as a therapy in VM and potentially in other disease states where cardiac function is equally compromised.

Earlier research by Barker and colleagues has demonstrated the interaction of SPARC with integrin-beta1 resulting in increased contractile signalling in lung fibroblasts through activation of ILK. Using SPARC null and WT cells they showed that SPARC is required for fibronectin-induced ILK-activation, which resulted in increased contractile signalling through MLC phosphorylation in these pulmonary fibroblasts [18]. In cardiomyocytes, MLC2v has been identified to be a critical regulator of cardiomyocyte contraction, by promoting actin-myosin interaction [27]. Our current working hypothesis is that SPARC increases cardiomyocyte contraction through its interaction with the integrin-beta1-ILK complex at the cardiomyocyte membrane. As a consequence, MLC phosphatase activity decreases intracellularly and hence ultimately increases the phosphorylation of MLC2v, causing increased actin-myosin interaction and thus augmented cardiomyocyte contraction (Figure 4). In line with Barker *et al.* [18] we have demonstrated an inotropic function of SPARC but this time in cardiomyocytes. SPARC increases cardiomyocyte fractional shortening possibly through its interaction with integrin-beta1 and increased downstream ILK signalling. Further studies will be required, however, to fully elucidate this mechanism.

**Figure 4.**
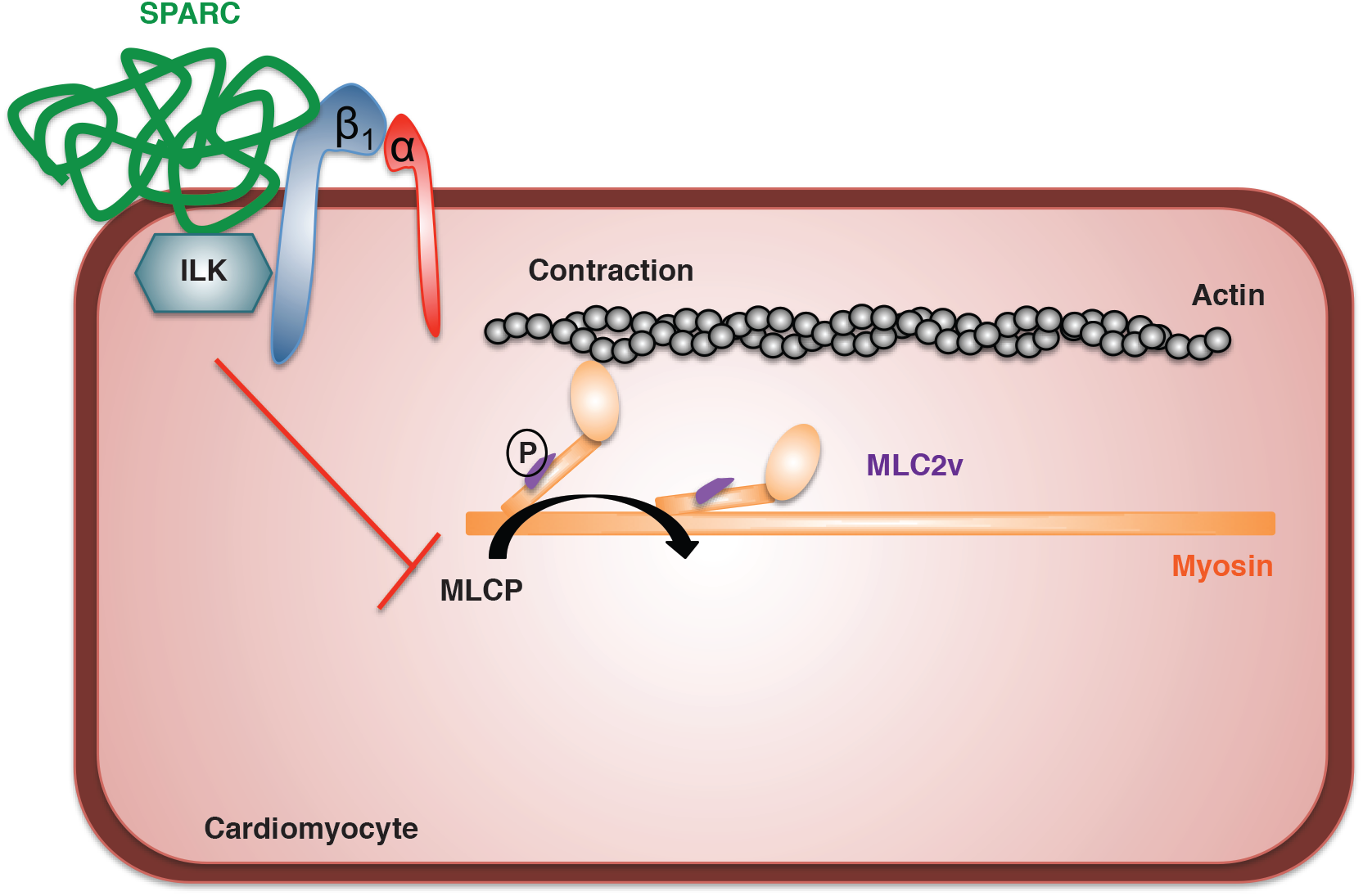
Proposed mechanism on how SPARC improves cardiomyocyte contraction. Our working hypothesis is that SPARC interacts with integrin-beta 1 and ILK on the cardiomyocyte membrane. This results in increased ILK signaling, blocking myosin light chain phosphatase (MLCP), and in this way increasing MLC phosphorylation and thus contraction.

Interestingly, in a study using monoclonal antibodies and peptides, the copper-binding domain of SPARC was identified to be required for the interaction of SPARC with integrin-beta1, resulting in increased ILK signalling. In the latter study, stressed lens epithelial cells displayed improved survival *in vitro* due to this interaction [28]. Mooney and colleagues also demonstrated improved survival through intergrin-beta1 signalling in mesangial cells, however SPARC failed to promote survival in this model.[29]

In the present study we did not investigate a potential protective effect of SPARC on myocyte survival, however we could not find evidence for decreased stress in VM hearts of SPARC overexpressing or SPARC treated mice, as was shown by equal amounts of fibrosis and CD45 positive cells, and the absence of cardiomyocyte hypertrophy in both subsets of hearts. Furthermore, SPARC overexpression did not result in altered levels of phosphorylated Akt, which is known to regulate cardiomyocyte hypertrophy and apoptosis [30, 31]. On the other hand, we did see a slight reduction in leukocyte infiltration in SPARC overexpressing hearts, 1 week after CVB3 infection. So if SPARC would provide any protective effect during VM, it is most likely through affecting leukocyte infiltration into the heart, and not by directly promoting cardiomyocyte survival.

Furthermore, we also demonstrate a rapid effect of SPARC on collagen cross-linking *in vivo* as collagen cross-linking is augmented in VM hearts with severe fibrosis, but not in VM hearts with little fibrosis or in healthy hearts, after 3 days of SPARC administration. Importantly, FS in the heart and of the isolated cardiomyocytes was higher in all animals which were administered SPARC.

## Conclusions

In conclusion, this study is the first to demonstrate a novel inotropic function for SPARC in the healthy heart, possibly by interacting with the integrin-beta1 on the cardiomyocyte membrane and resulting in altered downstream contractile signaling. Moreover, we have demonstrated the benefit of SPARC on contractile forces after coxsackie virus induced cardiac injury, which emphasizes the potential therapeutic application of this agent under these conditions, and perhaps provides proof of concept that this protein could also be of therapeutic benefit in other cardiac diseases where contractile function is diminished.

## Funding

This work was supported by a CARIM-funded PhD grant, and by the European Commission’s grants FIBROTARGETS [602904], MEDIA [261409], and ARENA [CVON 2011]. DMJ was funded by a postdoctoral fellowship from the FWO.

## Supporting Information

**Supplementary Figure 1.**
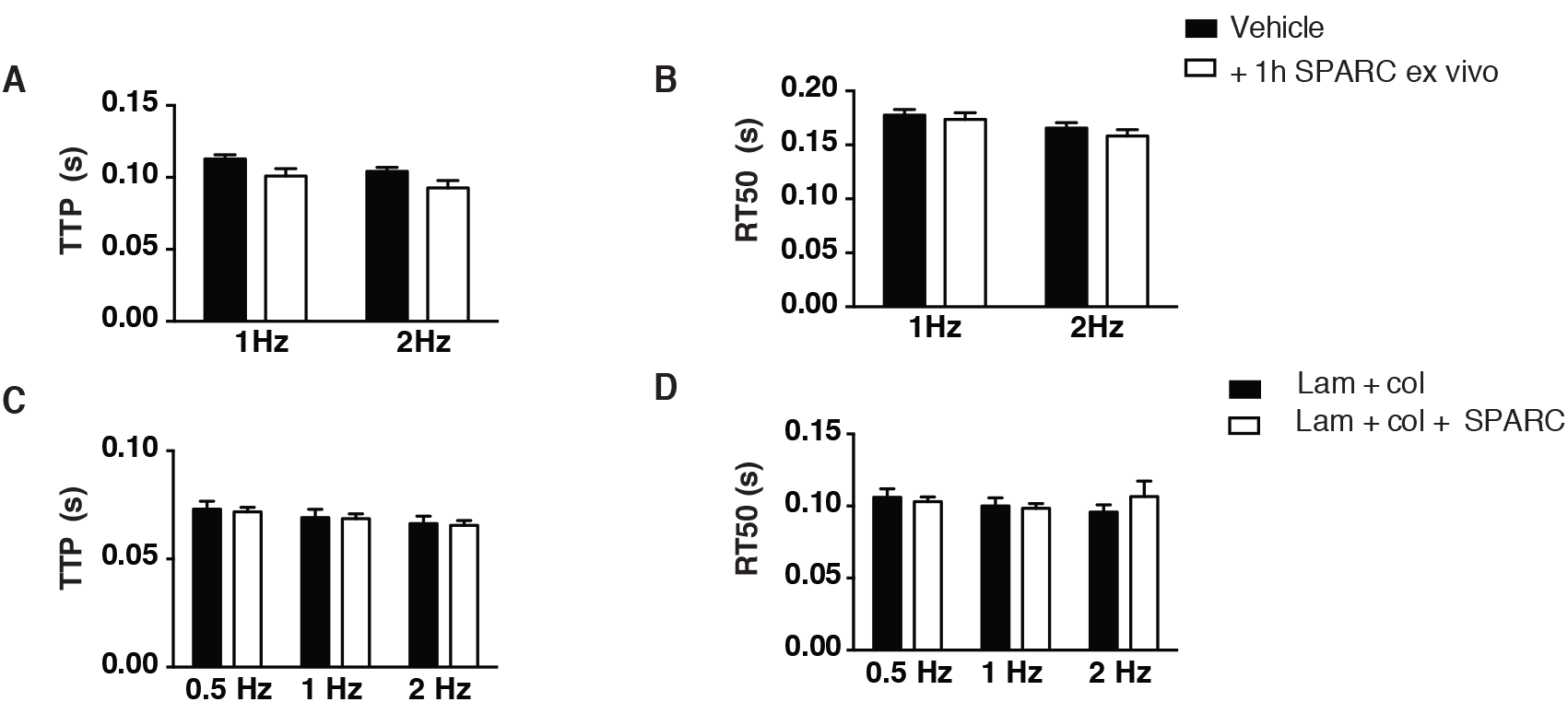
A,B Incubation of isolated adult mouse cardiomyocytes with recombinant SPARC for 1h *ex vivo* does not affect contraction – and relaxation times (TTP and RT50). C, D TTP and RT50 are not altered in rat cardiomyocytes grown on a matrix with SPARC. A,B N=4 mice and >4 cells per mouse, C,D N = 3 rats and >20 cells per rat

**Supplementary Figure 2.**
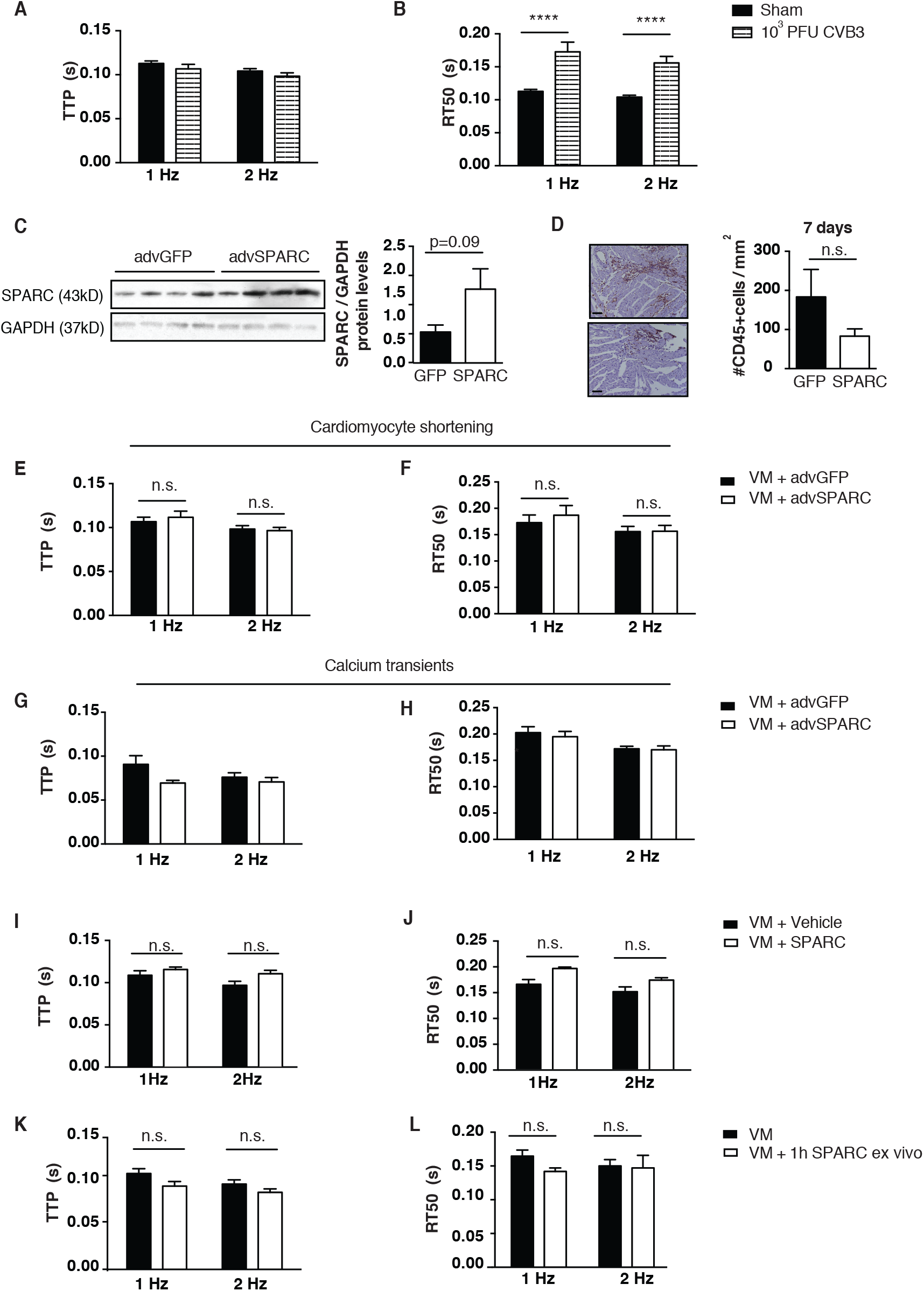
A, B Viral infection does not influence TTP but increases RT50 in isolated cardiomyocytes from virus-infected mice. C cardiac SPARC is almost significantly overexpressed in the adenoviral-SPARC injected group when compared to the control adenoviral-GFP injected mice, as shown by Western Blotting. D Slightly decreased cardiac inflammation, as measured by the amount of CD45 positive cells, was seen in the SPARC overexpressing group. E, F No effect on contraction or relaxation times was observed when SPARC was overexpressed. G,H There were no differences in the Ca^2+^ transient peak TTP or RT50 in cells from the SPARC overexpressing VM mice. I, J Cardiomyocytes from SPARC-treated mice demonstrated no differences in TTP or RT50. K,L When cardiomyocytes were isolated from severely sick, untreated mice, incubation of the cells with SPARC for 1h *ex vivo* did not influence TTP or RT50. A,B n= 11 for sham and n=13 for VM and >3 cells per mouse, C n=4 for both groups, D n=5 for both groups, E-H n=12 for advGFP group and n= 11 for advSPARC group and >3cells per mouse, I,J n=6 for VM+vehicle and n=7 for VM+SPARC, K,L n=13 for both groups and >3cells per mouse

